# Session duration affects the magnitude of post-exercise hypervolemia but not the erythropoietin response to acute high-intensity interval exercise

**DOI:** 10.1101/2024.09.03.611105

**Authors:** Thomas R. Tripp, Allison M. Caswell, Brittany A. Edgett, Martin J. MacInnis

**Affiliations:** Faculty of Kinesiology, University of Calgary, Calgary, Alberta, Canada

**Keywords:** Erythropoiesis, high-intensity interval training, plasma volume, exercise prescription

## Abstract

The increase in plasma volume ∼24 hours post-exercise may act as an erythropoietic signal, but this mechanism’s responsiveness to different exercise prescription variables is poorly understood. The purpose of this study was to determine the impact of high-intensity interval exercise duration on plasma volume and related responses. On separate days, 16 healthy, recreationally active participants (n=8 males; n=8 females) performed four (4×4) or eight intervals (8×4) consisting of 4 min at 105% critical power with 3 min recovery. Venous blood samples collected before, immediately after, and 24 hours after each HIIT session were used to measure hemoglobin concentration and hematocrit to calculate plasma volume changes. Erythropoietic and plasma volume regulating hormone concentrations were measured using ELISA kits. Plasma volume decreased immediately after both protocols (4×4: −4.4±3.5%, p<0.05; 8×4: −4.4±3.6%, p<0.05) but was only significantly elevated above baseline 24 hours after the 8×4 protocol (4×4: +1.0±7.1%, p>0.05; 8×4: +5.6±4.6%, p<0.05). Erythropoietin concentration ([EPO]) was higher than baseline 24 hours after the HIIT protocols (4×4: Pre vs 24 h post: 6.5±3.1 vs. 7.1±3.3 mIU/mL; 8×4: 6.9±3.7 vs. 7.3±3.7 mIU/mL; main effect of time, p<0.05) with no difference between protocols (p>0.05). [Aldosterone] was elevated immediately post-exercise after both protocols (4×4: Pre vs 0 h post: 295±151 vs. 544±259 pg/mL; 8×4: 335±235 vs. 821±553 pg/mL), but the 8×4 protocol caused a larger increase (interaction effect, p<0.05). That post-exercise hypervolemia may be exercise duration-dependent but is not required for increases in circulating EPO has important implications for endurance training aiming to increase oxygen delivery to active tissues.

**NEWS AND NOTEWORTHY:** This study is the first to show that both a common HIIT protocol length (4 x 4 min) and an extended HIIT protocol (8 x 4 min) similarly increased [erythropoietin] 24 h after exercise, despite only the extended protocol transiently increasing plasma volume. Previous works have investigated plasma volume regulation following different HIIT durations, but not explored links to erythropoietic signalling. This study’s findings have relevance for understanding the physiology of exercise-induced erythropoiesis.

## INTRODUCTION

Endurance training leads to blood volume expansion (1) and while the exact mechanisms are unresolved, exercise-induced changes in plasma volume are likely involved. Red blood cell proliferation is triggered by increased circulating erythropoietin concentration ([EPO]) in response to physiological and/or environmental stimuli, including hypoxia, exercise, and whole-body tilting (2), among others (3). Due to sweating and fluid shifting out of the vasculature, there is a decrease in blood volume during and immediately after exercise (4) that activates fluid retention hormone signalling pathways, including the renin-angiotensin-aldosterone system (RAAS). As a result, compensatory exercise-induced hypervolemia occurs ∼24 h after the exercise session, transiently decreasing hematocrit (5, 6). Over an extended period of repeated training, increased erythropoiesis gradually restores (approximately) the setpoint hematocrit with the sustained increase in plasma volume. The overall result is an increase in red blood cell volume, hemoglobin mass, and therefore total oxygen carrying capacity compared to pre-training (7). As total blood volume is an important determinant of maximal oxygen uptake (8) and submaximal exercise thresholds (9), understanding the influence of exercise prescription variables on the acute adaptive signalling pathways that augment blood volume adaptations is useful for optimizing adaptations to training.

As exercise duration determines the time during which fluid can be lost in response to exercise, it may be an important upstream determinant of fluid regulation and central venous pressure mechanisms implicated in exercise-induced erythropoiesis (2, 7). A recent study compared plasma volume following four, six, and eight high-intensity intervals, and showed similar increases in plasma volume (∼4-6%) 24 h after each duration (10). Those results suggested a limited role for duration, which is surprising given that exercise-induced hypervolemia and the concentrations of fluid regulating hormones are higher following more intense exercise (6, 11, 12). These contradictory findings–exercise intensity but not exercise duration affecting hypervolemia– necessitate a deeper investigation into the mechanisms connecting exercise duration, hypervolemia, and erythropoiesis. While previous studies have independently interrogated the plasma volume response to different exercise durations (10), or the potential underlying erythropoietic signalling pathways (2), to our knowledge, no study has simultaneously investigated the interaction of these mechanistic steps.

Our aim was to determine the role of exercise duration on the magnitude of post-exercise hypervolemia while examining potential upstream (plasma volume regulating hormone concentrations) and downstream signalling effects (erythropoietic hormone concentrations) in response to a common HIIT protocol (four 4-min intervals interspersed with 3 min of recovery) (13). We addressed this aim by doubling the session length from four intervals to eight intervals and monitoring the changes in plasma volume post-exercise. We hypothesized that doubling the volume of the HIIT session would increase, albeit not double, post-exercise hypervolaemia, based on augmented post-exercise hypervolemia when exercise volume is augmented by increasing exercise intensity (12). As previous research showing that plasma-volume regulating hormones are responsive to different exercise volumes via changes in exercise intensity (11, 12), we hypothesized that doubling the exercise volume would augment hormone responses (i.e., higher or lower, depending on the function of the hormone as a diuretic/anti-diuretic, respectively) and that the downstream erythropoietic responses would be similarly augmented.

## MATERIALS AND METHODS

### Participants

We recruited 16 healthy young males (n=8) and females (n=8) across a range of fitness levels to participate in this study (Table 1). Before any testing, a researcher explained the study purpose and all testing procedures to the participants. Participants provided written informed consent and completed a physical activity screening questionnaire (Get Active Questionnaire, Canadian Society for Exercise Physiology). This study received institutional ethics approval from the University of Calgary Health Research Ethics Board (REB22-0501) and complied with the Declaration of Helsinki, except for registration in a database.

**Table 1:** Participant characteristics.

| Variable | Pooled | Male | Female | Comparison<br>(ES, p-value) |
| --- | --- | --- | --- | --- |
| Number | 16 | 8 | 8 | N/A |
| Age (y) | 25 ± 5 | 26 ± 6 | 24 ± 4 | 0.10, 0.522 |
| Height (m) | 1.71 ± 0.09 | 1.77 ± 0.08 | 1.66 ± 0.08 | 0.52, 0.010 |
| Body Mass (kg) | 69.0 ± 10.5 | 75.3 ± 10.1 | 62.6 ± 6.4 | 0.45, 0.009 |
| BMI (kg/m<sup>2</sup>) | 23.5 ± 2.1 | 24.1 ± 2.7 | 22.8 ± 1.1 | 0.17, 0.246 |
| Body Fat (%) | 22.3 ± 7.7 | 16.4 ± 4.9 | 28.2 ± 5.0 | -0.82, <0.001 |
| VO<sub>2</sub>max (ml/kg<br>BM/min) | 46.1 ± 6.7 | 49.3 ± 5.8 | 42.9 ± 6.1 | 0.38, 0.050 |
| VO<sub>2</sub>max (ml/kg<br>FFM/min) | 58.7 ± 7.0 | 58.7 ± 7.3 | 58.7 ± 7.2 | 0.00, 0.996 |

ES, effect size (Cohen’s d); BMI, body mass index; BM, body mass; FFM, fat-free mass.

### Experimental Design

This study was part of a larger investigation examining the physiological determinants of critical power (9) and the effects of exercise duration on performance fatiguability (14). This manuscript only describes the procedures related to the research questions identified above. This study involved 8-9 laboratory visits separated into three distinct phases, occurring over the course of 4-5 weeks.

#### Phase 1: Baseline physiological testing

On their first and second visits to the laboratory, participants completed a ramp incremental test to exhaustion on a cycle ergometer to determine maximal oxygen uptake (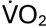max) and a dual-energy x-ray absorptiometry (DXA) scan to quantify fat-free mass, respectively.

#### Phase 2: Critical power determination

Participants completed 3-5 constant load cycling trials to task failure to determine critical power, which was used to prescribe the exercise intensity during the HIIT sessions in phase 3. Each trial was performed on a separate visit to the laboratory at a similar time of day (± 1 h), with a minimum of 48 h between trials.

#### Phase 3: High-intensity interval training protocols

In a randomized order one week apart, participants completed a HIIT session consisting of either 4 or 8 intervals. Blood samples were collected prior to each HIIT session, as well as immediately post-exercise and 24 h post-exercise to measure plasma volume changes, volume regulating hormone concentrations, and erythropoietic signals.

During this phase of the study, participants were instructed to control and record their fluid intake from ∼18 h prior to each HIIT session until the 24 h post-exercise blood draw. Fluid intake prescription is shown in Table 2. Participants recorded any additional fluid intake during their first experimental period and were asked to replicate their exact fluid intake during the second experimental period. Additionally, we asked participants to avoid caffeine and alcohol from 18 h prior to each HIIT session until the 24 h post-exercise blood draw.

**Table 2:** Fluid intake control schedule.

| Time Span | Fluid Intake Prescription |
| --- | --- |
| ~Dinner to sleep (day before exercise) | 15 ml/kg |
| Wake-up to exercise start | 15 ml/kg |
| Post-exercise until dinner | Volume equivalent to weight lost to sweat (1 L per kg), plus habitual daily intake* |
| Dinner to sleep | 15 ml/kg |
| Wake-up to 24 post-exercise blood draw | 15 ml/kg |

\* For whichever trial was performed second, participants consumed the same habitual fluid volume during this period as they had for the first trial. The only difference was the volume consumed to balance the weight lost due to sweat.

### Data Collection

#### Ramp incremental exercise test

Participants completed a ramp incremental exercise test on an electromagnetically braked cycle ergometer (Velotron; Racermate Inc., Seattle, WA, USA) to quantify 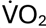max. After a baseline data collection period of 4 min at 50 W, the power output increased linearly at a rate of 25 W/min (1 W every 2.4 s). Participants were allowed to see their cadence but were blinded to elapsed time and power output throughout the test. Participants were encouraged to cycle until exhaustion and researchers provided strong, verbal encouragement throughout the test. Along with volitional task failure, the test was terminated when participants could no longer keep their cadence above 60 rpm. Heart rate was monitored using a chest strap (Polar H10; Polar Electro, Kempele, Finland). Gas exchange was monitored using a two-way non-rebreathing valve (Hans Rudolph Inc., Shawnee KS, USA) connected to a metabolic cart via a mixing chamber (Quark CPET, Cosmed, Rome, Italy). 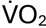max was calculated as the highest 30 s 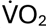 achieved during the test. The final power output from the ramp test at volitional exhaustion was taken as peak power output (PPO).

#### Dual-energy x-ray absorptiometry

Participants underwent a DXA scan in a Lunar iDXA device (General Electric Healthcare, Chicago, IL, USA). The scan was used to quantify fat-free mass and body fat percentage.

#### Critical power testing and calculation

Each constant-load cycling trial consisted of 5 min baseline cycling at 50 W and then a step transition to the trial power output. Participants were allowed to see their cadence throughout the trial but were blinded to the power output and elapsed time. Participants were instructed to cycle as long as possible and were given strong verbal encouragement to continue cycling. Volitional exhaustion or an inability to maintain a cadence above 60 rpm were used to determine task failure. Power outputs were selected pseudo-randomly based on percentages of PPO from the ramp incremental test with the aim of obtaining three trials lasting between 2 and 15 min. Critical power was calculated using three equations: linear work-time, hyperbolic power-time, and linear power-inverse time. The equation that gave the lowest standard error of the critical power estimate was selected.

#### High-intensity interval exercise protocol

The high intensity interval exercise sessions consisted of 4 min at 105% of critical power (severe intensity domain) interspersed with 3 min at 15% of critical power. In a randomized order one week apart, participants completed one trial of four intervals (4×4) and one trial of eight intervals (8×4). The average interval power output during each session was 202 ± 46 W. Participants completed 3.77 ± 0.82 kJ of total work during the 4×4 HIIT sessions and 7.34 ± 1.63 kJ during the 8×4 HIIT session. Heart rate, rating of perceived exertion (RPE; 6-20 top to bottom; Borg, 1982), and rating of general fatigue (fatigue; 0-10 bottom to top; Micklewright et al., 2017) were recorded during each HIIT trial. Average heart rate during interval 4 was slightly higher during the 8×4 HIIT session (8×4: 172 ± 11 vs. 4×4: 168 ± 13 bpm; paired t test: p = 0.002) but RPE (median [range] – 8×4: 15 [11-19] vs. 4×4: 15 [10-19]; Wilcoxon signed-rank test: p = 0.998) and fatigue (8×4: 6 [3-8] vs. 4×4: 5 [2-8]; Wilcoxon signed-rank test: p = 0.383) were not significantly different.

Participants were weighed (without shoes but wearing their exercising clothes) on a balance beam scale (Detecto-Medic; Detecto Scales Inc., Brooklyn, NY, USA) before and immediately after each exercise bout to calculate body mass lost to sweat (17) and to estimate the minimum fluid intake requirement following each exercise bout (Table 2).

#### Blood sampling

Blood was collected from an antecubital vein using a 21G butterfly needle connected to a tube holder. Three tubes (3-5 mL) were drawn in the following order: serum, heparin, EDTA. Serum samples were allowed to clot at room temperature for 30 minutes prior to centrifugation at 3000 x g for 15 min at 4 °C. Heparin and EDTA-treated plasma samples were immediately centrifuged at 3000 x g for 15 min at 4 °C. Serum and plasma samples were transferred to 1.5 ml microcentrifuge tubes and stored at −30 °C until analysis.

After the final tube was collected, the tube holder was removed, and a heparinized syringe was used to collect ∼1 mL of blood for measurement of hemoglobin and hematocrit (ABL80; Radiometer, Copenhagen, Denmark). These values were used to calculate the change in plasma volume from baseline to each post-exercise time point based on the equation of Dill and Costill (1974).

#### ELISA experiments

Concentrations of renin ([renin]), N-terminal proatrial natriuretic peptide ([NT-proANP]), and EPO were measured in EDTA plasma samples using commercially available ELISA kits (Human Renin DuoSet ELISA: DY4090; Human NT-proANP DuoSet ELISA: DY8247-05; Human Erythropoietin/EPO DuoSet ELISA: DY286-05; R&D Systems, Inc., Minneapolis, MN, USA). Serum [aldosterone] was measured using a commercially available ELISA kit (Aldosterone ELISA Kit (Colorimetric): KA1883; R&D Systems, Inc.). For measurement of insulin-like growth factor 1 concentration ([IGF-1]), serum samples were pretreated with ethanol and HCl, per manufacturer suggestion and following the protocol of Daughaday et al. (1980) and measured using a commercially available ELISA kit (Human IGF-I/IGF-1 DuoSet ELISA, DY291; R&D Systems, Inc.). All samples were assayed in duplicate and final concentrations were adjusted based on the change in plasma volume measured from baseline (18, 20). For example, if plasma volume increased by 5% (effectively lowering the concentration assuming no change in the hormone mass), hormone concentrations would be multiplied by 1.05 to account for the dilutionary effect.

Prior to any sample analysis, we performed antibody grid experiments to optimize capture and detection antibody concentrations and spike-recovery/linearity experiments to optimize sample preparation/dilution (21). These experiments were performed for all analyte kits other than aldosterone, which was validated by the manufacturer.

### Statistical Analysis

Data were checked for normality with Shapiro-Wilk tests and visual inspection of the Q-Q plots, equality of variance using Levene’s test, and sphericity using Greenhouse-Geisser epsilon. Data were checked for outliers using the robust regression and outlier removal (ROUT) method at Q = 1%. We removed one participant from both the [renin] and [NT-proANP] analysis and a second participant from the [NT-proANP] analysis due to extremely high values.

Post-exercise plasma volume responses (as a percentage change from baseline) and blood hormone concentrations were assessed using a two-way repeated measures ANOVA with the within-subject factors of time (Pre [hormone concentrations only], 0 h, and 24 h) and session duration (4×4 or 8×4 HIIT). A significant ANOVA result was followed with Tukey’s multiple comparisons tests to identify means that were significantly different. As the plasma volume calculations did not permit statistical comparisons with a true baseline value, the mean plasma volume change in each condition, at each time point (i.e., four groups), was compared to a value of zero using one-sample t tests.

## RESULTS

### HIIT session intensity measures: heart rate, RPE, and fatigue

Average heart rate across the entire session, average heart rate during the intervals, and peak heart rate were all higher during the 8×4 compared to the 4×4 HIIT protocol (Table 3; p < 0.001 for each). RPE and ratings of fatigue during the final interval were both higher during the 8×4 compared to the 4×4 HIIT protocol (Table 3, p < 0.001 for each).

**Figure 1:**
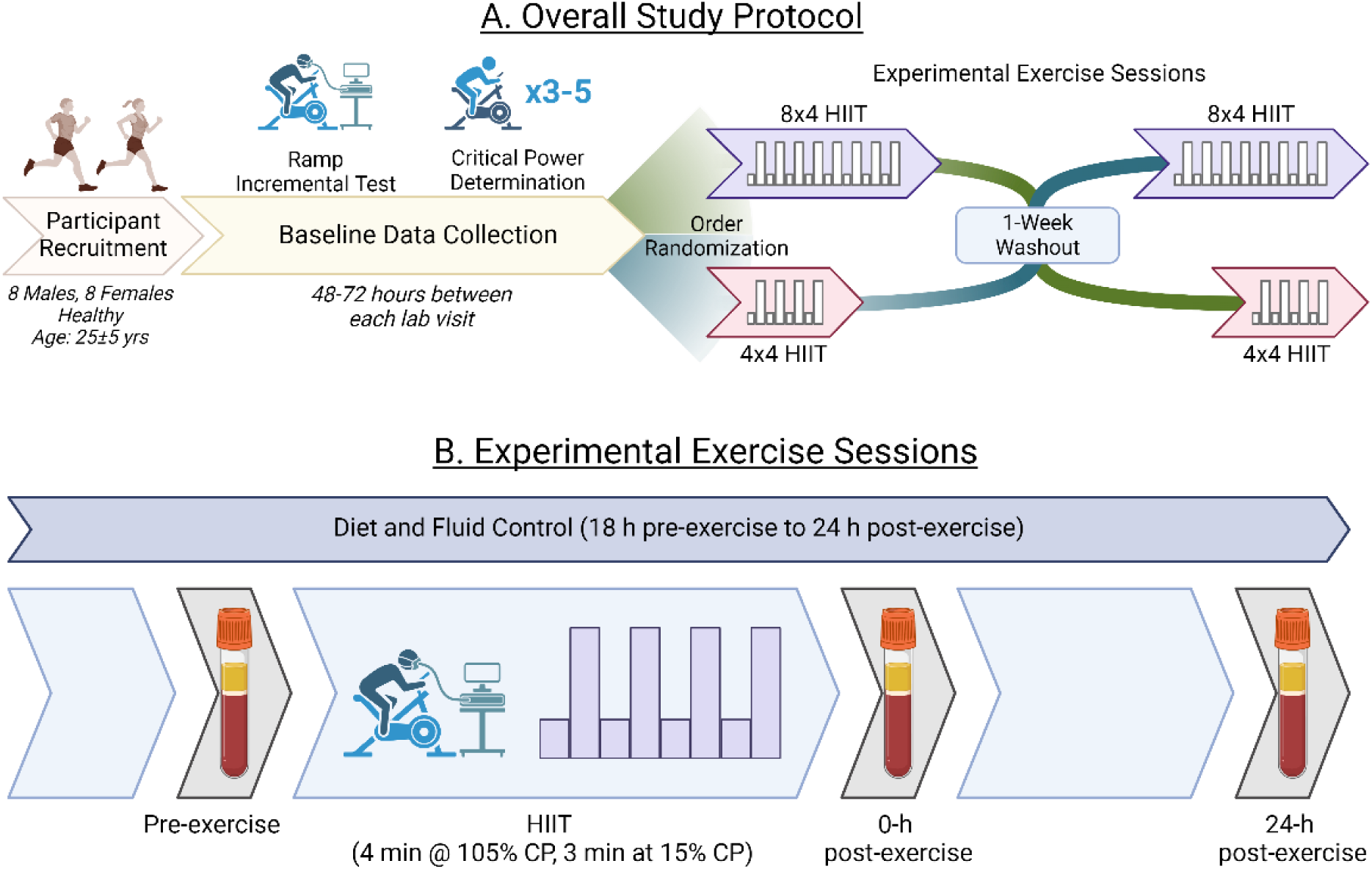
Protocol schematic. The overall study timeline is shown in panel A. After initial participant screening, which involved a maximal oxygen uptake (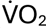max) test and dual-energy x-ray absorptiometry (DXA) scan, we determined critical power to use in prescribing the high-intensity interval training (HIIT) sessions. In a randomized, counterbalanced order, participants completed 4×4 and 8×4 HIIT sessions with venous blood collected before, immediately after, and 24 hours after each session (B). The detailed timeline for the experimental sessions is shown in panel B. Created with BioRender.com.

**Table 3:** High-intensity interval training session characteristics.

| Variable | 4x4 Session | 8x4 Session | Comparison (p-value) |
| --- | --- | --- | --- |
| Session Average HR<br>(bpm) | 143 ± 13 | 153 ± 11 | < 0.001 |
| Interval Average HR<br>(bpm) | 162 ± 13 | 169 ± 9 | < 0.001 |
| Peak HR<br>(bpm) | 177 ± 12 | 183 ± 10 | < 0.001 |
| Final Interval RPE<br>(6-20) | 15 (10-19) | 16.5 (13-20) | < 0.001 |
| Final Interval Fatigue<br>(0-10) | 5 (2-8) | 7 (5-10) | < 0.001 |

RPE and Fatigue data are shown as median (range). HR, heart rate; RPE, rating of perceived exertion; Fatigue; rating of general fatigue.

### Plasma volume responses to HIIT

Plasma volume decreased significantly immediately following both HIIT protocols, with no effect of session duration (4×4: −4.4 ± 3.5 vs. 8×4: −4.4 ± 3.6%; p = 0.999; Figure 2A). The 8×4 HIIT protocol (+5.6 ± 4.6%; p < 0.001) caused a significant increase in plasma volume 24 hours post-exercise that was not observed with the 4×4 HIIT protocol (+1.1 ± 7.1%; p = 0.564). Body weight also decreased immediately following exercise (4×4: Pre: 69.4 ± 11.1 vs. 0 h: 69.1 ± 11.1 kg; 8×4: 69.7 ± 11.2 vs. 68.9 ± 11.1; Figure 2B), with a larger decline following the 8×4 protocol (−1.1 ± 0.2%) compared to the 4×4 protocol (−0.5 ± 0.2%; p < 0.001).

**Figure 2:**
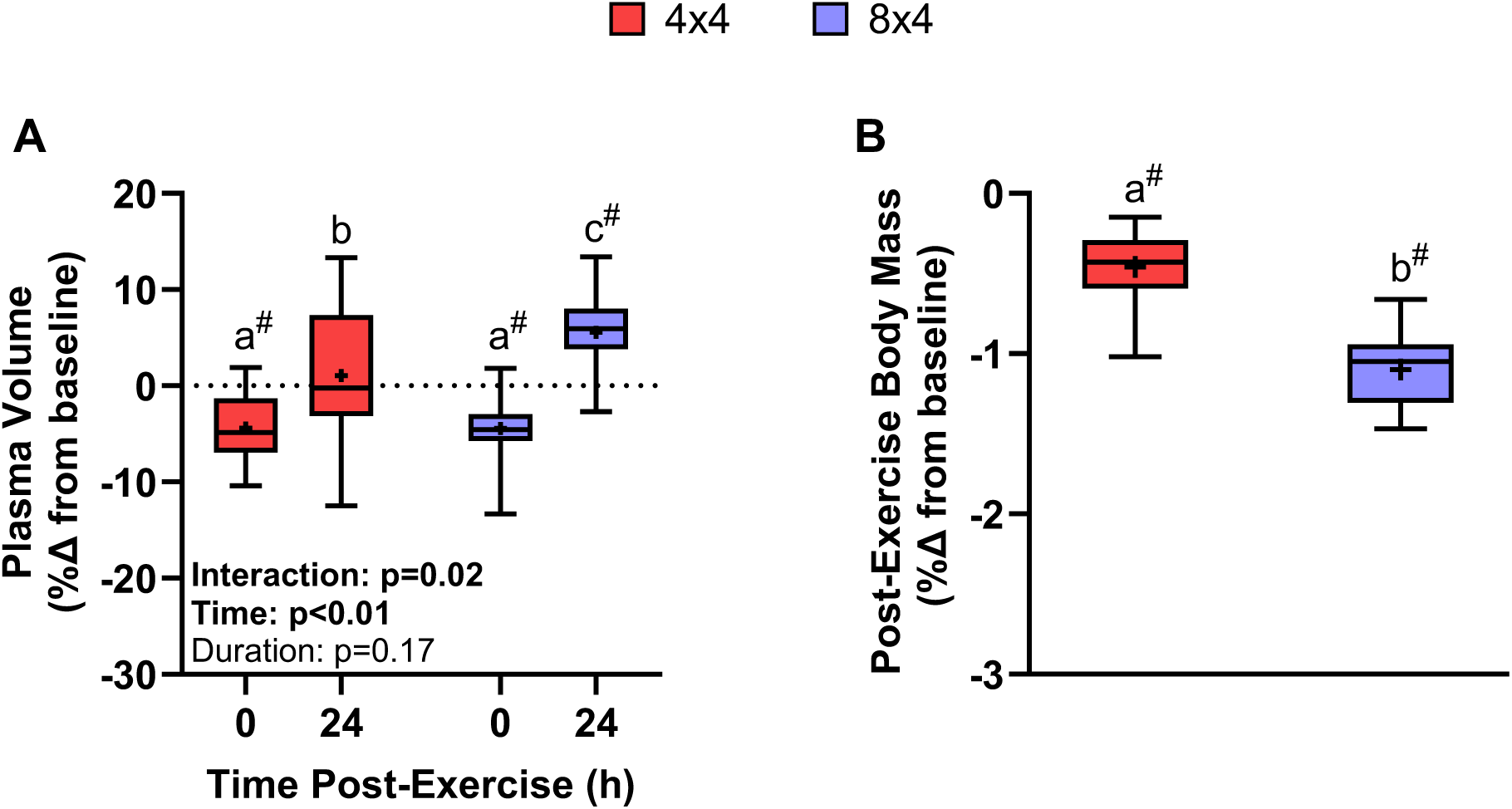
Plasma volume and body mass responses to two high-intensity interval training (HIIT) durations. Plasma volume decreased significantly immediately following both exercise bouts (A). Plasma volume returned to baseline values 24 hours after the 4×4 HIIT session but remained elevated above baseline 24 hours after the 8×4 HIIT session. Body mass declined significantly from baseline following each HIIT session, with a significantly larger decline after the 8×4 HIIT session (B). Plots depict interquartile range (box), with median (line), mean (cross), and error bars showing the range from minimum to maximum. Two-way repeated measures ANOVA results are shown with significant effects bolded. Points with different letters are significantly different from one another (Sidak’s post-hoc tests). ^#^ Significantly different from 0 (p < 0.05, one-sample t test). n = 16.

### Plasma volume regulating hormones

Plasma [renin] was significantly elevated from Pre at 0 h post-exercise (Figure 3A, main effect: Pre: 500 ± 195 vs. 0 h: 590 ± 218 pg/ml; p = 0.002) following both 4×4 (Pre: 500 ± 189 vs. 0 h: 562 ± 192 pg/ml) and 8×4 HIIT (501 ± 207 vs. 619 ± 244). Plasma [renin] at 24 h post-exercise (main effect: 532 ± 236; 4×4: 545 ± 260; 8×4: 519 ± 217 pg/ml) was not different from Pre (p = 0.378).

**Figure 3:**
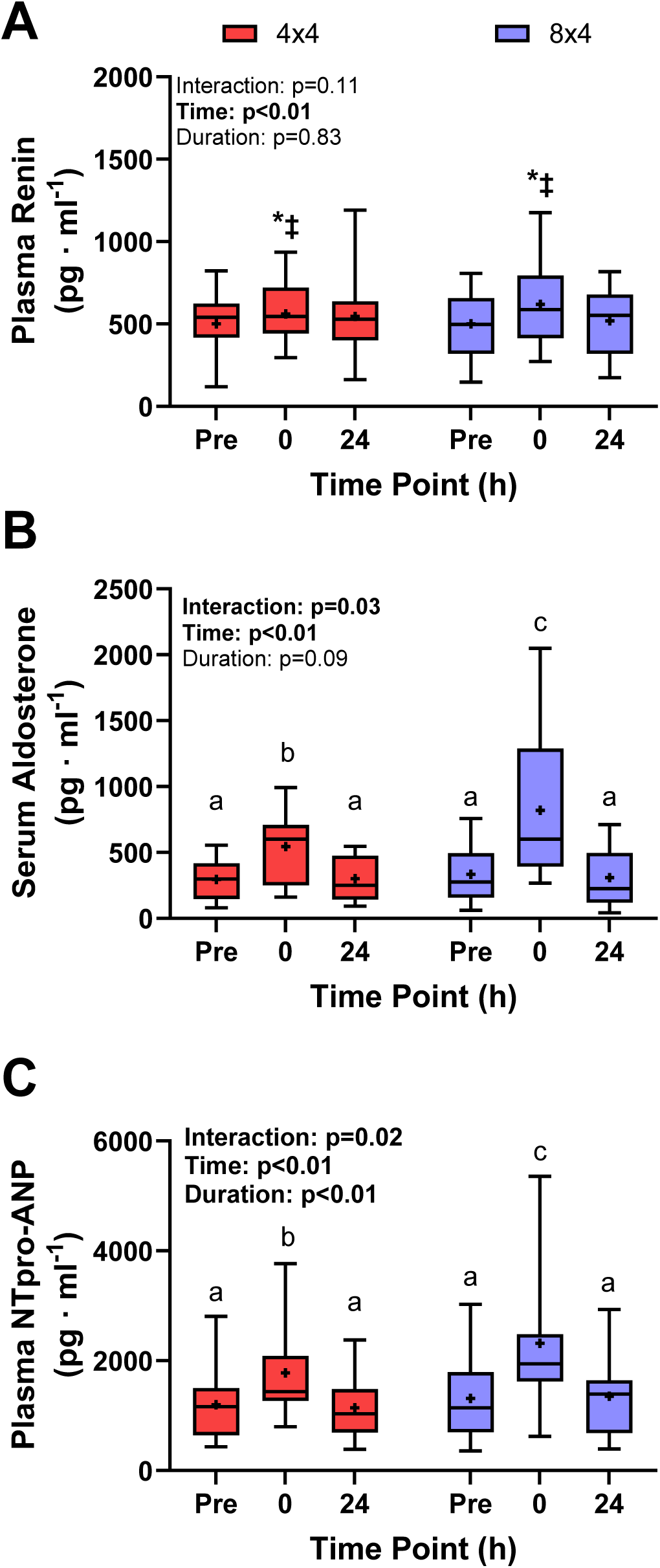
Plasma-volume regulating hormone concentrations following 4×4 and 8×8 high-intensity interval training (HIIT) sessions. Changes in hormones relating to plasma volume regulation were examined in plasma and serum. Plasma renin concentrations (A) were elevated immediately post-exercise and returned to baseline values by 24 hrs post-exercise in both conditions. For serum aldosterone (B) and plasma N-terminal proatrial natriuretic peptide (NT-proANP) (C), the exercise-induced increase in concentration was greater following 8×4 HIIT than 4×4 HIIT. Plots depict interquartile range (box), with median (line), mean (cross), and error bars showing the range from minimum to maximum. Two-way repeated measures ANOVA results are shown with significant effects bolded. When a significant interaction is present, points with different letters are significantly different from one another (Sidak’s post-hoc tests). Symbols indicate statistically significant differences between time points: * different from Pre, ‡ different from 24. n = 15, 16, and 14 for renin, aldosterone, and NT-proANP, respectively.

Serum [aldosterone] was increased by both HIIT protocols, but the increase at 0 h post-exercise was greater following 8×4 HIIT (Pre: 335 ± 235 vs. 0 h: 821 ± 553 pg/ml, p < 0.001; Figure 3B) than 4×4 HIIT (4×4: 295 ± 151 vs. 545 ± 259 pg/ml, p = 0.006). Serum [aldosterone] decreased to baseline values by 24 h post-exercise in both conditions (4×4: 300 ± 162, p = 0.998; 8×4: 310 ± 231 pg/ml, p = 0.941).

Plasma [NT-proANP] showed a significant interaction between time and HIIT duration (Figure 3C, p = 0.020), with a significant increase from Pre at 0 h post-4×4 HIIT (Pre: 1196 ± 628 vs. 0 h: 1779 ± 881 pg/ml; p < 0.001) but a larger increase at 0 h post-8×4 HIIT (Pre: 1310 ± 713 vs. 0 h: 2320 ± 1382 pg/ml, p < 0.001). 24 h following each condition, plasma [NT-proANP] returned to baseline values (4×4: 1144 ± 566, p = 0.872; 8×4: 1354 ± 775 pg/ml, p = 0.909).

### Erythropoietic hormones

Plasma [EPO] was unchanged from Pre at 0 h post-exercise (Figure 4A, main effect means: Pre: 6.7 ± 3.4 vs.0 h: 6.4 ± 3.3 mIU/ml, p = 0.384; 4×4: 6.5 ± 3.1 vs. 6.2 ± 3.3; 8×4: 6.9 ± 3.7 vs. 6.7 ± 3.4), but plasma [EPO] was increased 24 h post-exercise compared to Pre (main effect: 7.2 ± 3.4, p = 0.024; 4×4: 7.1 ± 3.3; 8×4: 7.3 ± 3.7 mIU/ml). Serum [IGF-1] remained unchanged following the 4×4 (Pre: 237 ± 82, 0 h: 238 ± 82, 24 h: 244 ± 86 pg/ml; Figure 4B) and the 8×4 HIIT (Pre: 229 ± 85, 0 h: 230 ± 82, 24 h: 234 ± 79).

**Figure 4:**
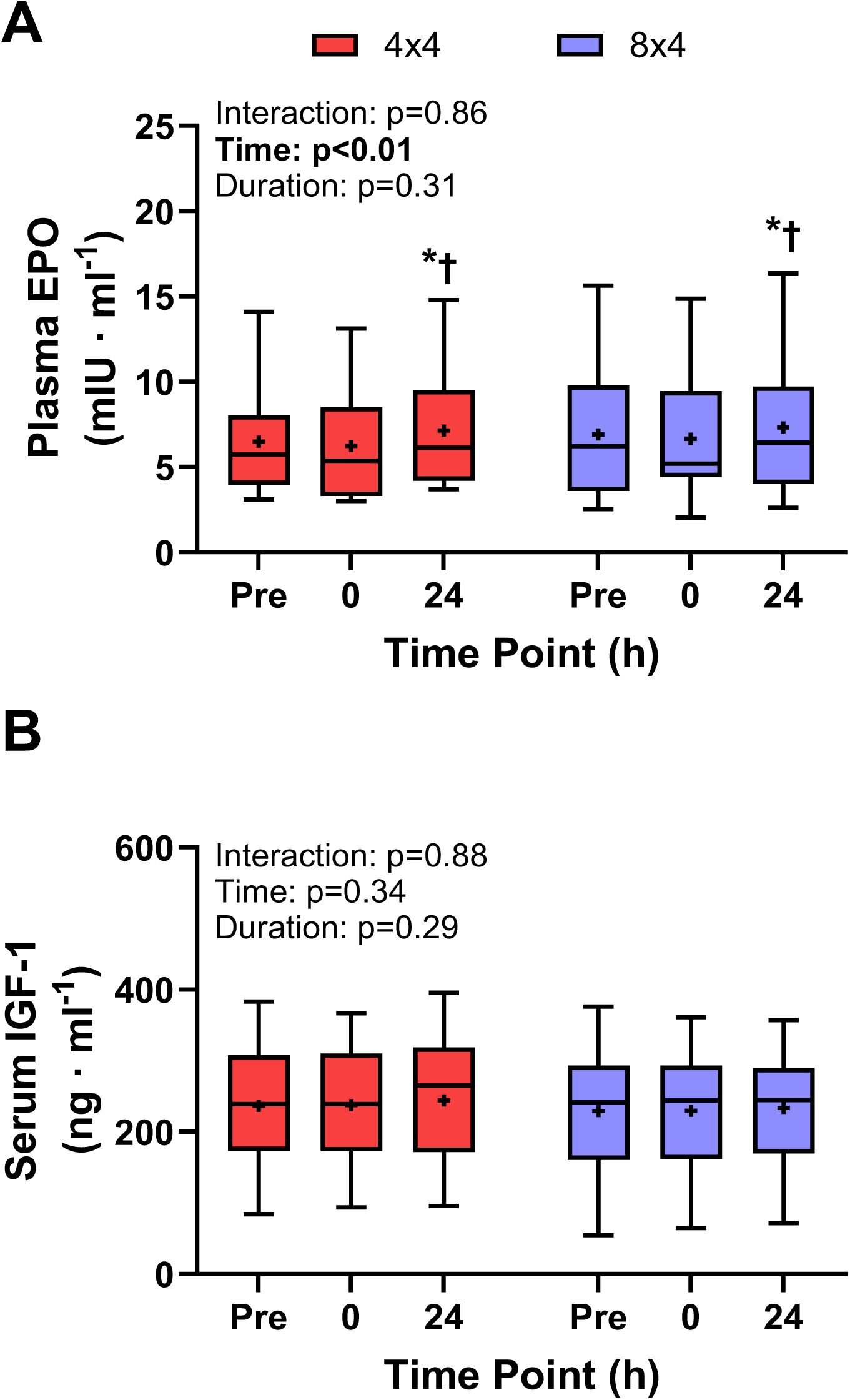
Erythropoietic hormone concentrations following 4×4 and 8×4 high-intensity interval training (HIIT) sessions. Changes in erythropoietin (EPO; A) and insulin-like growth factor 1 (IGF-1; (B) were examined in plasma and serum. IGF-1 remained unchanged following exercise. EPO was significantly elevated 24 hrs post-exercise. Plots depict interquartile range (box), with median (line), mean (cross), and error bars showing the range from minimum to maximum. Two-way repeated measures ANOVA results are shown with significant effects bolded. Symbols indicate statistically significant differences between time points: * different from Pre, † different from 0. n = 16.

## DISCUSSION

This study aimed to determine the role of exercise duration in post-exercise hypervolemia by comparing plasma volume responses to two different durations of HIIT: a common 4 x 4 min HIIT protocol and an extended 8 x 4 min HIIT protocol. To gain a greater mechanistic insight, we also measured the concentrations of several plasma volume regulating and erythropoietic hormones to understand the upstream (i.e., determinants of plasma volume changes) and downstream (i.e., signalling effects of plasma volume changes) implications of our observed post-exercise hypervolemia results. Regarding our primary aim, we found post-exercise hypervolemia following the longer (8×4 min) but not the shorter (4×4 min) HIIT protocol. The main novel finding of the present study was that the increase in plasma [EPO] was similar between low- and high-volume HIIT despite greater post-exercise hypervolemia and plasma volume regulating hormone signalling responses after the extended-duration HIIT.

### Plasma volume pattern following HIIT protocols

The 8 x 4 min HIIT protocol caused an increase of ∼5% in plasma volume ∼24 hours following exercise. As hypothesized, the longer exercise duration resulted in a greater hypervolemic response, likely because of greater reductions in body mass (i.e., sweat loss) and activation of fluid retention mechanisms in the kidneys. This difference in fluid retention was not related to the magnitude of plasma volume lost during the exercise, as both protocols caused a similar decrease in plasma volume measured immediately following the bouts (approximately −4.5%). The magnitude of hypervolemia seemed to be related to the duration over which the body was exposed to the decreased plasma volume, rather than the magnitude of post-exercise hypovolemia. In agreement with this, previous work has shown that markedly different states of hypovolemia immediately after exercise (i.e., 50% different between intense intervals and continuous exercise) lead to similar hypervolemia ∼8-22 hours later (∼5% in both conditions; (22).

Given that the first half of the 8×4 protocol in our study was identical to the entire 4×4 protocol, participants must have completed the second half of the protocol in a state of hypovolemia. The ∼30-min of additional exercise time before the recovery process began allowed greater time to stimulate fluid-retention pathways. Plasma [renin] was significantly elevated following both protocols without a statistically significant difference between conditions, though it was numerically higher after the 8 x 4 min HIIT. Greater pathway stimulation is further supported by our findings that serum [aldosterone] (downstream of renin in the RAAS pathway) and plasma [NT-proANP] were increased by both protocols but to a greater extent immediately following the 8 x 4 min HIIT than the 4 x 4 min protocol. Greater [NT-proANP] may promote a more potent diuretic effect, but the presence of 24 h post-exercise hypervolemia indicates to us that the balance was in favour of fluid retention following the 8 x 4 min protocol.

Hypervolemia following 8 x 4 min HIIT agreed with previous reports. Several studies have shown that plasma volume increased between 4-10% (4, 10, 23, 24) after a similar exercise protocol (8 x 4 min at 85% 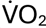max, with 5 min at 40% 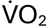max between each interval). The only differences from our protocol were the method of prescribing exercise intensity and the duration of the rest period, which do not appear to have affected the plasma volume outcomes.

Some major differences with previous literature are evident when examining the effect of exercise duration on post-exercise hypervolemia. Two studies have shown similar hypervolemic responses to different exercise durations (10, 25). One of these studies found no statistically significant hypervolemia following three durations of continuous exercise (30, 60, and 90 mins at 50% peak cycling power output), which may have been due to insufficient exercise intensity or statistical power (25). Specific to HIIT, Nelson et al. (ref 10) found a 6.6% increase in plasma volume following 4 x 4 min HIIT that was not significantly greater after 6 x 4 min or 8 x 4 min. In contrast, we found no change on average after this same duration (16 min of high-intensity cycling), with post-exercise hypervolemia only evident following 8 x 4 min HIIT. Both studies included similar participant cohorts; controls for diet, fluid intake, and exercise order; and were performed at similar absolute intensities (214 ± 42 W in Nelson et al. compared to 202 ± 46 W in our study). Our contradictory finding is surprising and challenges the notion that exercise duration is unimportant (or is unimportant beyond four intervals) for triggering post-exercise hypervolemia in response to HIIT. Taken with our body mass data (larger decrease after higher volume) and mechanistic hormone data (larger increases after higher volume), we suggest that exercise duration is a critical factor determining the hypervolemia response to a single exercise session but acknowledge that further research is necessary to clarify disagreements between studies.

### Erythropoietic signalling

Plasma volume expansion has inherent benefits to exercise performance (26), but our primary interest in plasma volume expansion was as an early signalling event for eventual blood volume and hemoglobin mass expansion. Post-exercise hypervolemia may decrease oxygen availability to various tissues due to hemodilution, and EPO is a critical signalling hormone released in response to tissue hypoxia (27). We saw an increase in [EPO] 24 hours following both exercise sessions but no change in [IGF-1], which is known to promote erythroid cell proliferation and differentiation (28). IGF-1 has been shown to increase immediately following brief, intense exercise (29), but this effect may have been caused by changes in plasma volume rather than IGF-1 release (18, 20). Indeed, the uncorrected [IGF-1] in our study was slightly higher immediately post-exercise (main effect means: Pre: 233 ± 82 vs. 0 h: 243 ± 82, p = 0.0133), an effect that disappeared upon correction (Figure 3B). The decoupling of hypervolemia and erythropoietic signalling responses to common- and extended-duration HIIT suggests to us that hypervolemia may be a sufficient stimulus for increasing [EPO] post-exercise, but circulating [EPO] can increase even when hypervolemia does not occur.

The contribution of other mechanisms and signalling pathways may be responsible for the increase in [EPO] present after the 4×4 HIIT in the absence of significant plasma volume expansion. The most notable similarity between the plasma volume responses of the two sessions was in the immediate post-exercise plasma volume, which decreased by ∼5%. Previous work has shown that plasmapheresis is capable of stimulating EPO release, with the authors suggesting increased renal O_2_ consumption (via sodium resorption) as the key factor to lower renal PO_2_ (30). Alternatively, other tissues may produce or release EPO in response to hypoxia-dependent and hypoxia-independent mechanisms (31, 32). For example, increased *Epo* gene expression has been shown in skeletal muscle following exercise (33). Whether one of these mechanisms was responsible for the similar increase across the different exercise durations is not clear and requires further investigation.

The single session was sufficient to generate a detectable change in [EPO], regardless of exercise duration/volume. Our results, showing increased RAAS signalling, post-exercise hypovolemia, and delayed hypervolemia, are consistent with the two potential fluid-retention mechanisms discussed above acting upstream of the EPO response (7, 30). Although we detected an increase in [EPO] 24 hours following HIIT, it is possible that the time points measured herein were not optimal for fully describing these changes; previous studies have shown elevated [EPO] between 1 hour (2) and 3 hours post-stimulus (34). Thus, it is possible that a difference in [EPO] between conditions occurred in this time window and was not detected in the present study. A more granular time course of changes in plasma volume (22) may help elucidate a potential role in post-exercise EPO stimulation.

The obvious implication of our results is that, based on the similar [EPO] response, the 4×4 HIIT protocol would be as effective for increasing blood volume as the 8×4 HIIT protocol. Evidence from previous training studies indicates that this outcome is possible, as protocols employing 4×4 HIIT regularly report improvements in 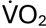max and exercise performance in a variety of populations (35–37); however, other studies directly measuring hematological adaptations suggest that greater exercise volume during a training protocol (albeit *via* increased exercise frequency rather than session duration) does augment blood volume expansion (38). An important difference between the present study and training studies is the repetition of the exercise session numerous times in close proximity to one another, providing the opportunity to augment the response (whether that be a hypervolemia-dependent or -independent response). As the present study was designed to investigate single HIIT sessions, differences in the magnitude of the hypervolemic response and its relevance to the downstream erythropoietic events could emerge (or disappear) with repeated exercise sessions as in regular training.

### Experimental considerations

We included male and female participants in our sample to improve the generalizability of our results. The study was powered to detect a difference between the exercise protocols rather than an interaction between duration and sex, or a main effect of sex, so we did not perform any sex-based analyses. Data disaggregated by sex are provided to allow others to plan studies investigating sex differences in this area (Supplemental Figure 1). The timing of our blood measurements was selected to balance time points most likely to capture post-exercise hypervolemia and to show changes in the hormones upstream and downstream of the phenomenon. It was also necessary to limit the total number of blood samples drawn to minimize participant burden and to avoid influencing any outcomes that may be sensitive to blood withdrawal. Between these considerations, we prioritized capturing the plasma volume changes, but may have missed the peak in acute [EPO] or [IGF-1] responses following exercise as a result.

## CONCLUSONS

This study investigated the impact of exercise duration on erythropoietic signalling and post-exercise hypervolemia by comparing plasma volume and hormone responses to two different HIIT protocols. Our findings revealed that doubling the exercise volume from four to eight intervals significantly increased plasma volume 24 hours post-exercise, suggesting that exercise duration plays a crucial role in stimulating fluid retention. Elevated concentrations of plasma volume regulating hormones following the longer HIIT protocol indicate a greater activation of the RAAS. Critically, we found similar increases in [EPO] following both protocols 24 hours post-exercise, showing a decoupling of hypervolemia and erythropoietic responses that suggests hypervolemia may be sufficient but not necessary for EPO stimulation. Further research is needed to elucidate the interplay between exercise prescription variables and adaptive signaling pathways to optimize endurance training outcomes.

---

**Supplemental Figure 1:**
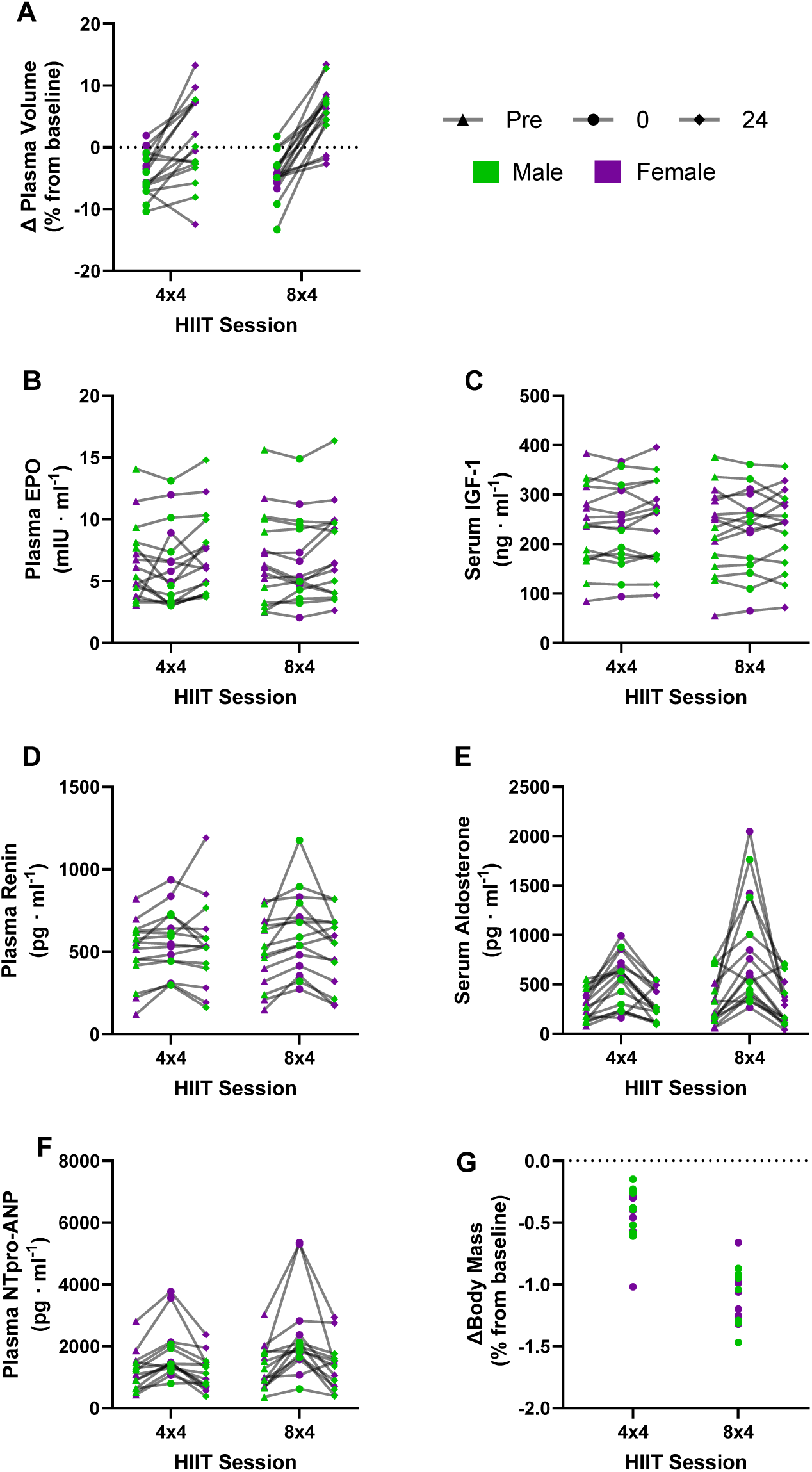
All variables showing individual data disaggregated by sex

